## Supplementary material for "Non-targeted isomer-sensitive N-glycome analysis reveals new layers of organ-specific diversity in mice": Supplmentary information

**Supplementary Data**

DeCon2 parameter file (“SampleParameterFile.xml”)

Found Mass List (“Annotated_MassList.csv”)

Semi-quantitative histograms of SNOG-filtered LC-MS data of all samples (“full N-glycome.bmp”)

Semi-quantitative histograms of Neu5Ac-associated eSNOG-filtered LC-MS data of all samples (“Neu5Ac N-glycome.bmp”)

Semi-quantitative histograms of Neu5Gc-associated eSNOG-filtered LC-MS data of all samples (“Neu5Gc N-glycome.bmp”)

Semi-quantitative histograms of Antennary-fucose-associated eSNOG-filtered LC-MS data of all samples (“Antennary Fuc N-glycome.bmp”)

Precursor-independent MS/MS-based profiling for Neu5Ac across all tissues (“MSMS_Neu5Ac.bmp”)

Precursor-independent MS/MS-based profiling for Neu5Gc across all tissues (“MSMS_Neu5Gc.bmp”)

Precursor-independent MS/MS-based profiling for fucose across all tissues (“MSMS_Fucose.bmp”)

**Supplementary Information**

Supplementary Note 1. Calculation of the SNOG-score

Supplementary Note 2. Calculation of the extended SNOG-scores (eSNOG-score)

Supplementary Figure 1. Precursor-independent MS/MS-based N-glycome profiling of 20 mouse tissues

Supplementary Figure 2. Precursor-independent MS/MS-based N-glycome profiling of 20 mouse tissues

Supplementary Figure 3. Impact of SNOG-score filtering on data quality

Supplementary Figure 4. Impact of SNOG-score filtering on the total ion current (TIC) and number of unique mass bins across all samples

Supplementary Figure 5. Impact of SNOG-score filtering on data quality of the liver 1 data set

Supplementary Figure 6. Empiric determination of the cut-off values for the extended SNOG-scores (eSNOG)

Supplementary Figure 7. Tissue specific expression of specific N-glycan modifications

Supplementary Figure 8. MS/MS supported identification of a N-glycan with the composition Hex_6_HexNAc_6_Fuc_4_GlcA_1_ carrying a fucosylated HNK-1 epitope in kidney

Supplementary Figure 9. Profiling the isomeric complexity of the murine N-glycome

Supplementary Figure 10. Profiling the isomeric complexity of the murine N-glycome

Supplementary Table 1. Overview over diagnostic fragment ions used for sub-structural profiling of the mouse N-glycome

**Supplementary Notes**

Supplementary Note 1. Calculation of the SNOG-score

The SNOG-score is a metric utilized to effectively differentiate signals originating from N-glycans from those originating from contaminants, such as polysaccharides. In experiments involving porous graphitic carbon (PGC)-LC-MS/MS, N-glycans are commonly reduced. The resulting reduced GlcNAc fragment ion (224.1 amu) serves as a diagnostic marker for N- (or O)-glycans. To compute the SNOG-score specific to mass bins (0.01 Da mass bins), the mean relative intensity of the 224.1 amu fragment ion is calculated across all MS/MS spectra within each precursor mass bin. This calculation is performed on a per-sample basis across all detected mass bins. Mass bins with a SNOG-score exceeding 0.03 are selected to construct a sample-specific target list, from which precursor-intensity information is retrieved.

Supplementary Note 2. Calculation of the extended SNOG-scores

The extended SNOG-scores (eSNOG) serve to categorize N-glycans based on sub-structural characteristics such as Neu5Ac-sialylation, antenna-fucosylation, or GlcNAc-sulfation. These scores are calculated in a similar manner to the original SNOG-score. Specifically, the mean relative intensity of one or more diagnostic fragments, which are specific to the modification of interest, is calculated across all MS/MS spectra within the corresponding precursor mass bin. This calculation is performed on a per-sample basis across all identified mass bins. The cutoff values for different eSNOG-scores vary depending on the specific modification and have been determined empirically (SI Table 1).

**Supplementary Figures**

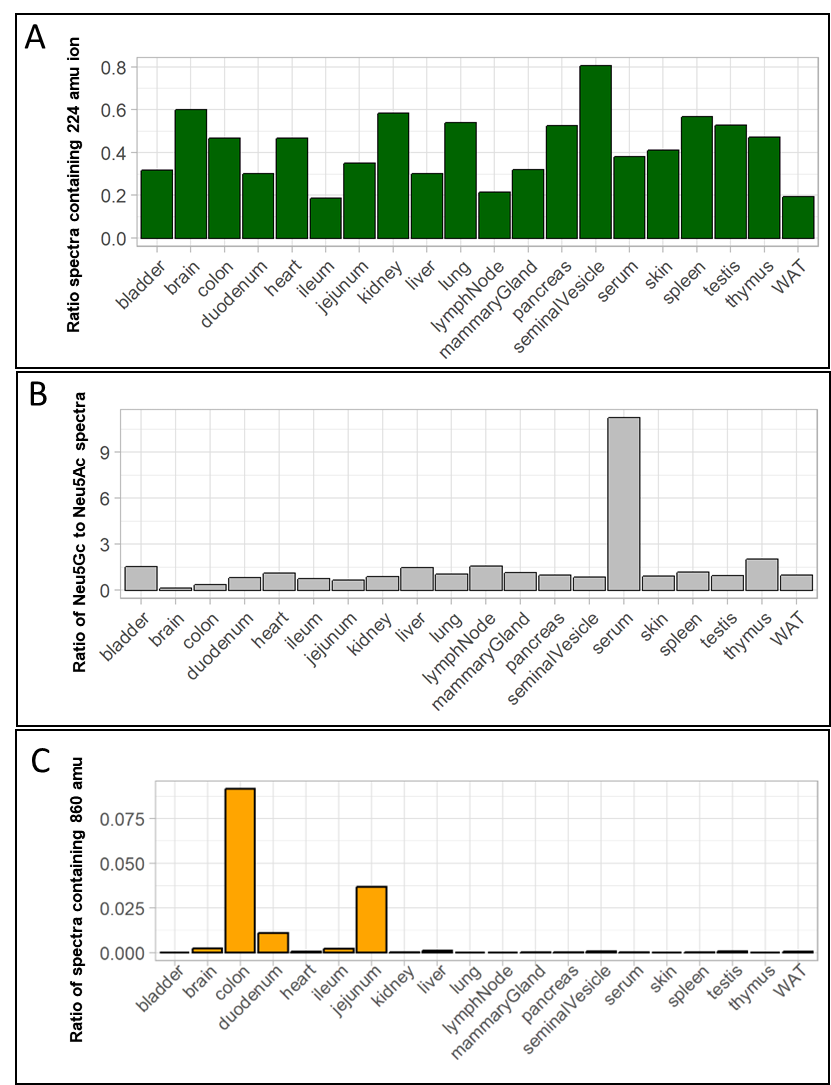

**SI Figure 1.** **Precursor-independent MS/MS-based N-glycome profiling of 20 mouse tissues. (A)** Relative abundance of MS/MS spectra containing the glycan-specific fragment ion 224.1 amu (reduced HexNAc). **(B**) Ratio of spectra containing the diagnostic fragment ion for Neu5Gc (290.1 amu) and spectra containing the diagnostic fragment ion for Neu5Ac (274.1 amu). **(C)** Ratio of spectra containing the fragment ion indicative for the Sda-antigen (860.3 amu). Only N-glycan associated spectra are considered (hence containing the fragment ion 224.1 amu). “WAT” - white adipose tissue.

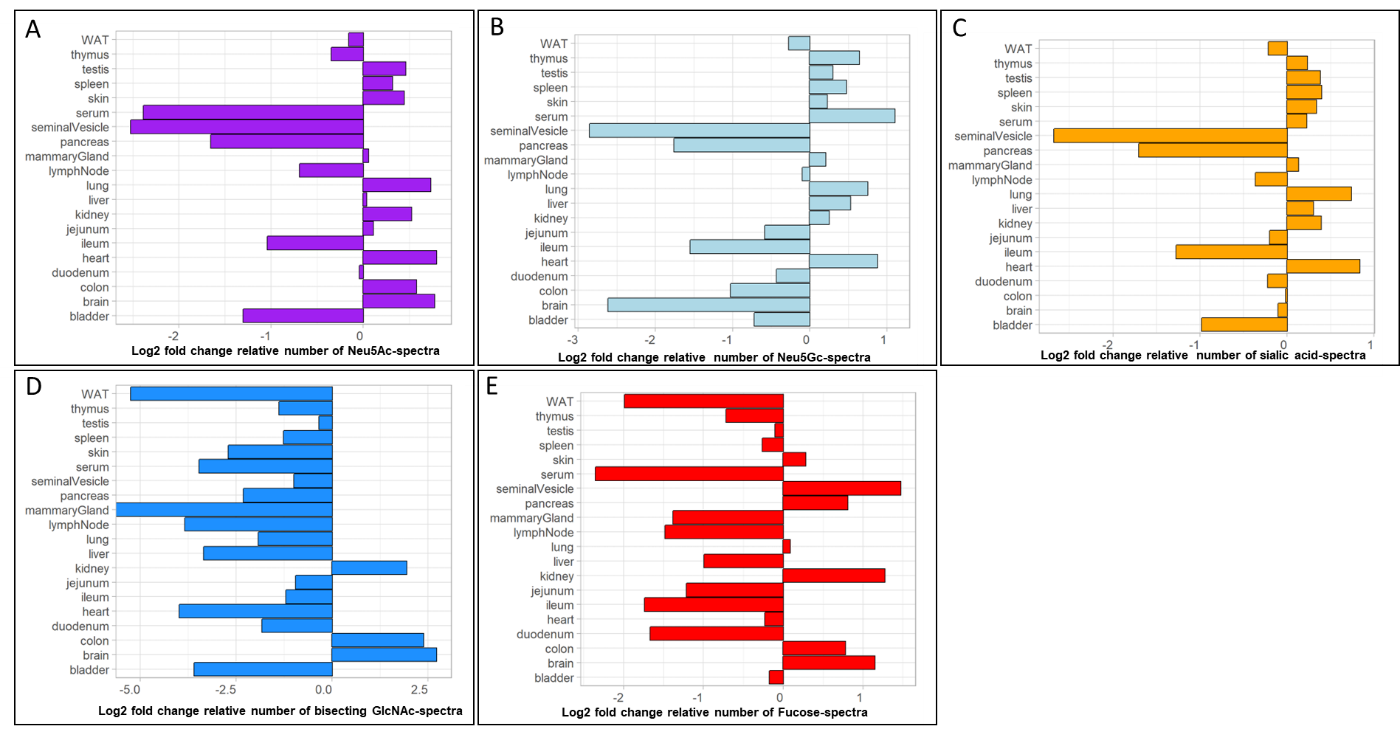

**SI Figure 2**. **Precursor-independent MS/MS-based N-glycome profiling of 20 mouse tissues**. The graphs show the log2-fold change of the number of MS/MS spectra containing a specific diagnostic fragment ions in relation to the mean across all tissues. **(A**) Log2-fold change of the number of MS/MS containing a Neu5Ac-specific fragment ion (274.1 amu) (B) Log2-fold change of the number of MS/MS containing a Neu5Gc-specific fragment ion (290.1 amu). **(C)** Log2-fold change of the number of MS/MS containing a Neu5Ac- or Neu5Gc-specific fragment ion (274.1 amu or 290.1 amu)**. (D)** Log2-fold change of the number of MS/MS containing a bisecting GlcNAc-specific fragment ion (792.3 amu). **(E)** Log2-fold change of the number of MS/MS containing a fucose-specific fragment ion (512.2 amu). Because of gas-phase re-arrangement of fucose-residues, this fragment ion does not discriminate between core- and distal fucose. Only N-glycan associated spectra are considered (hence containing the fragment ion 224.1 amu). “WAT” - white adipose tissue.

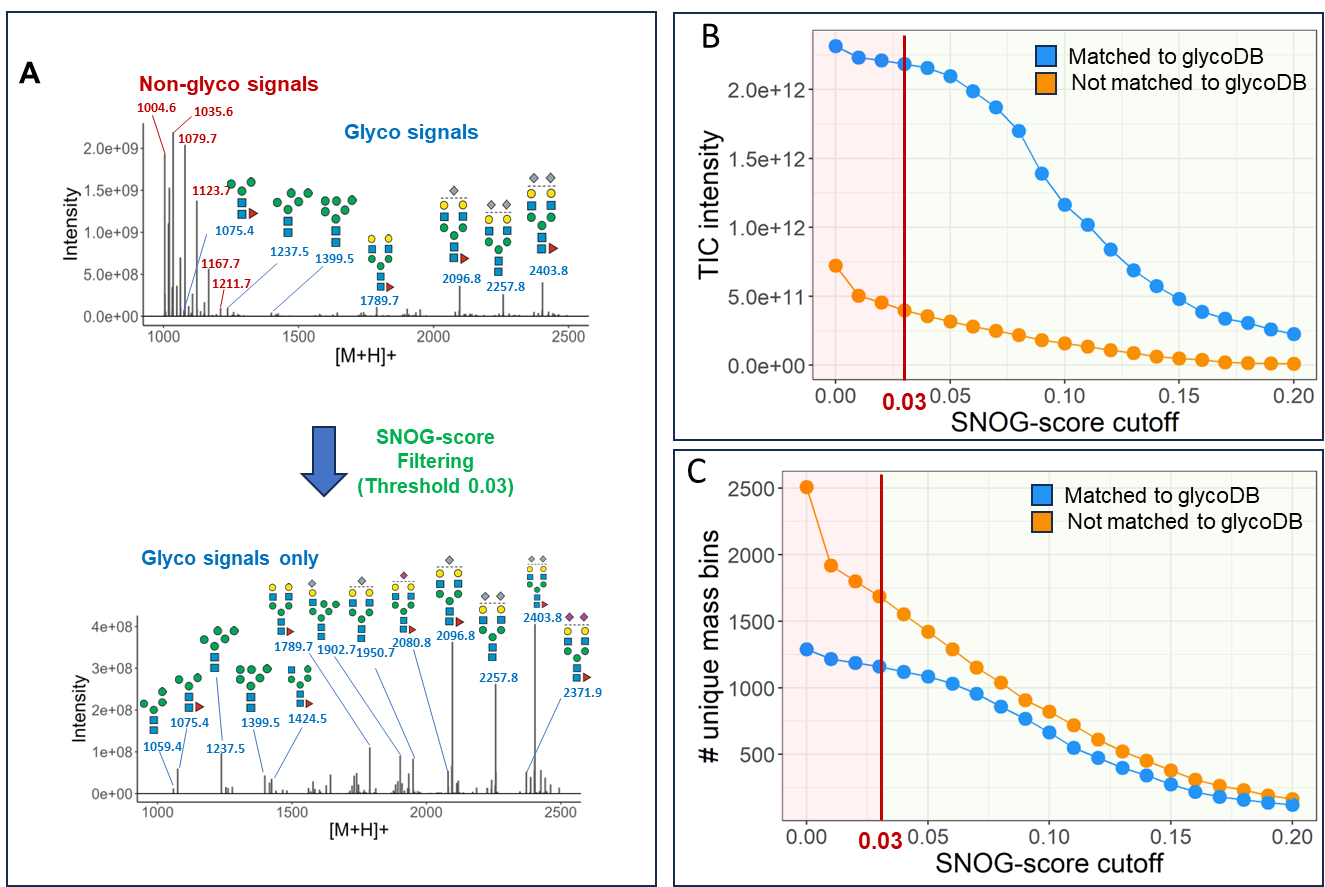

**SI Figure 3. Impact of SNOG-score filtering on data quality. (A)** SNOG-filtering of the lymph node 1 data set. Filtering effectively removes signals derived from contaminants and retains N-glycan associated signals. **(B)** Impact of different SNOG-score cutoff values on the accumulated total ion current (TIC) of all analyzed samples. **(C)** Impact of different SNOG-score cutoff values on the number of unique mass-bins across all analyzed samples. Blue dots represent values corresponding to mass bins which have an entry in our glycoDB (including sodium and ammonium adducts), orange dots represent values which have no entry in our glycoDB. N-glycan cartoons depict tentative structure assignments based on composition, using the „Symbol Nomenclature for Glycans“ (SNFG). “WAT” - white adipose tissue.

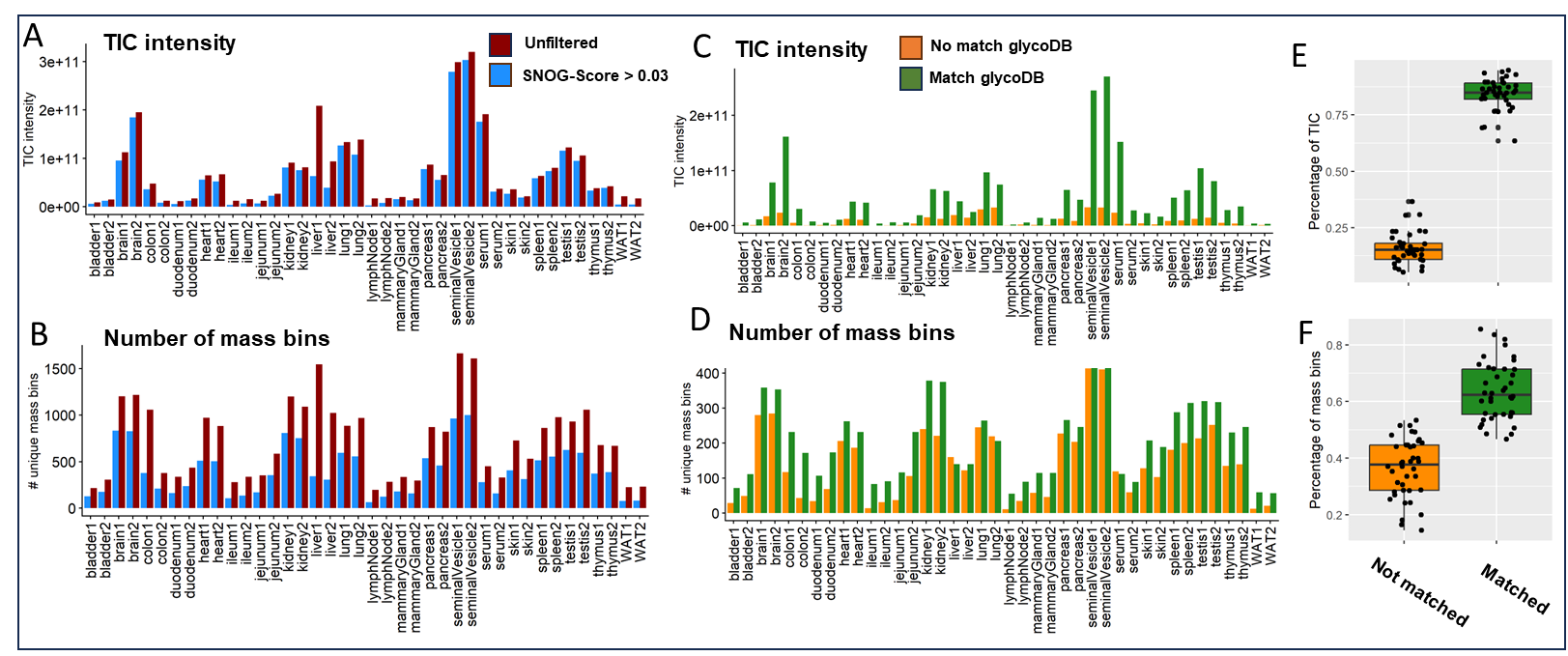

**SI Figure 4. Impact of SNOG-score filtering on the total ion current (TIC) and number of unique mass bins. (A)** Impact of the SNOG-score filtering with a cutoff at 0.03 on the TIC, stratified across all samples. Red bars represent the TIC of the unfiltered data, blue bars represent the filtered data. **(B**) Impact of the SNOG-score filtering with a cutoff at 0.03 on the number of unique mass bins, stratified across all samples. Red bars represent the TIC of the unfiltered data, blue bars represent the filtered data. **(C)** Distribution of the TIC of the SNOG-score filtered data set (> 0.03) categorized into mass bins that have an entry in our glycoDB (green) and into mass bins which have no entry in our glycoDB (including sodium and ammonium adducts (orange) across all samples**. (D)** Distribution of the number of unique mass bins of the SNOG-score filtered data set ( > 0.03) categorized into mass bins that have an entry in our glycoDB (green) and into mass bins which have no entry in our glycoDB (including sodium and ammonium adducts) (orange) across all samples.(**E)** Boxplots of the percentage compositions of the TIC derived from mass bins which have an entry in our glycoDB (including sodium and ammonium adducts) (green) or have no entry in our glycoDB (orange) across all samples.(**F)** Boxplots of the percentage compositions of the number of unique mass-bins which have an entry in our glycoDB (including sodium and ammonium adducts) (green) or have no entry in our glycoDB (orange) across all samples. “WAT” - white adipose tissue.

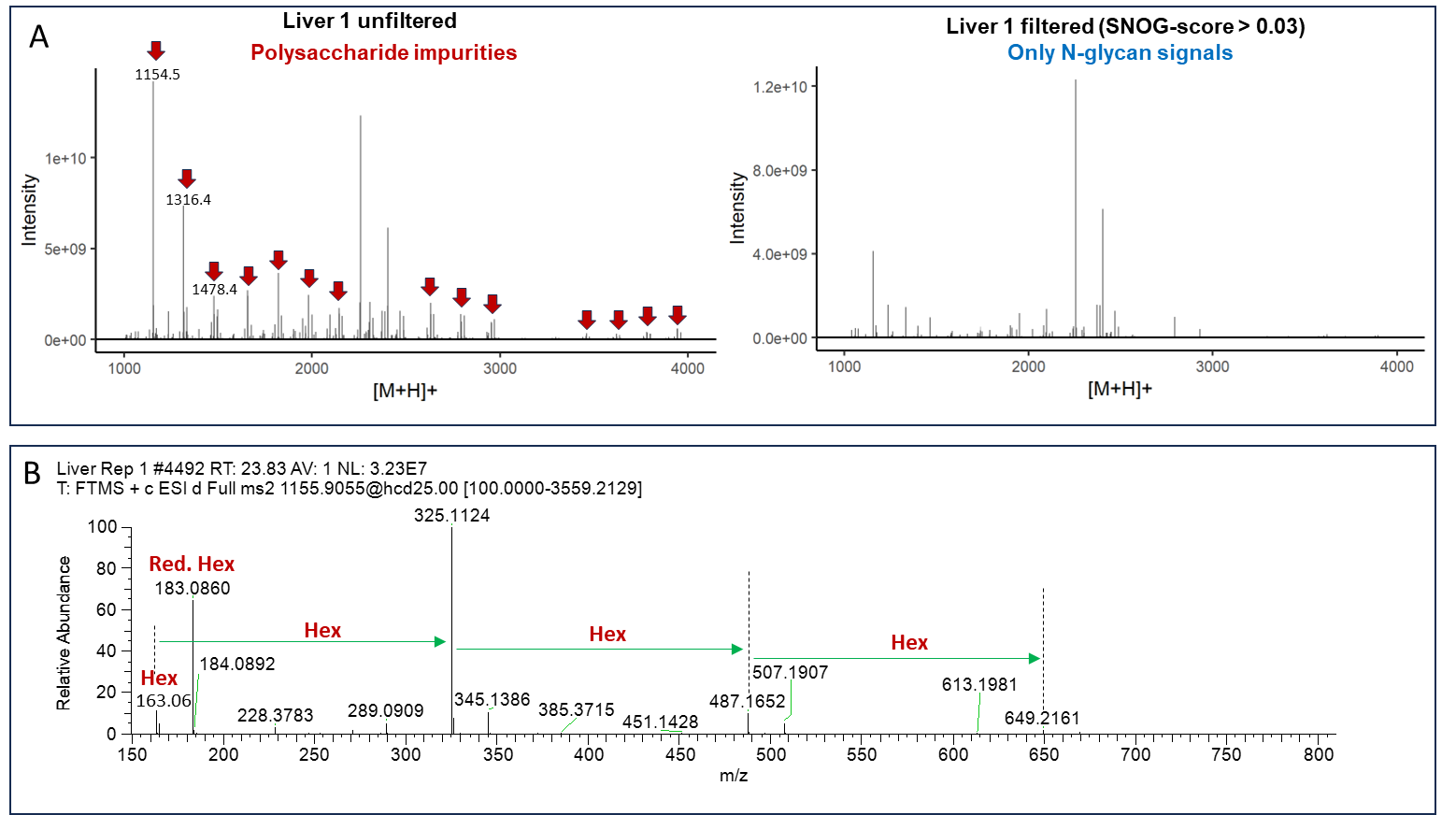

**SI Figure 5**. **Impact of SNOG-score filtering on data quality of the liver 1 data set. (A)** Comparison between the unfiltered (left) and filtered liver 1 data set (right). The red arrows depict signals derived from polysaccharide impurities commonly observed in N-glycan preparations. The SNOG-score filter efficiently removes impurity-derived signals. **(B**) Exemplary MS/MS spectrum of a polysaccharide impurity derived from the liver 1 data set. The absence of a 224.1 amu diagnostic fragment ion leads to the removal of the respective precursor mass.

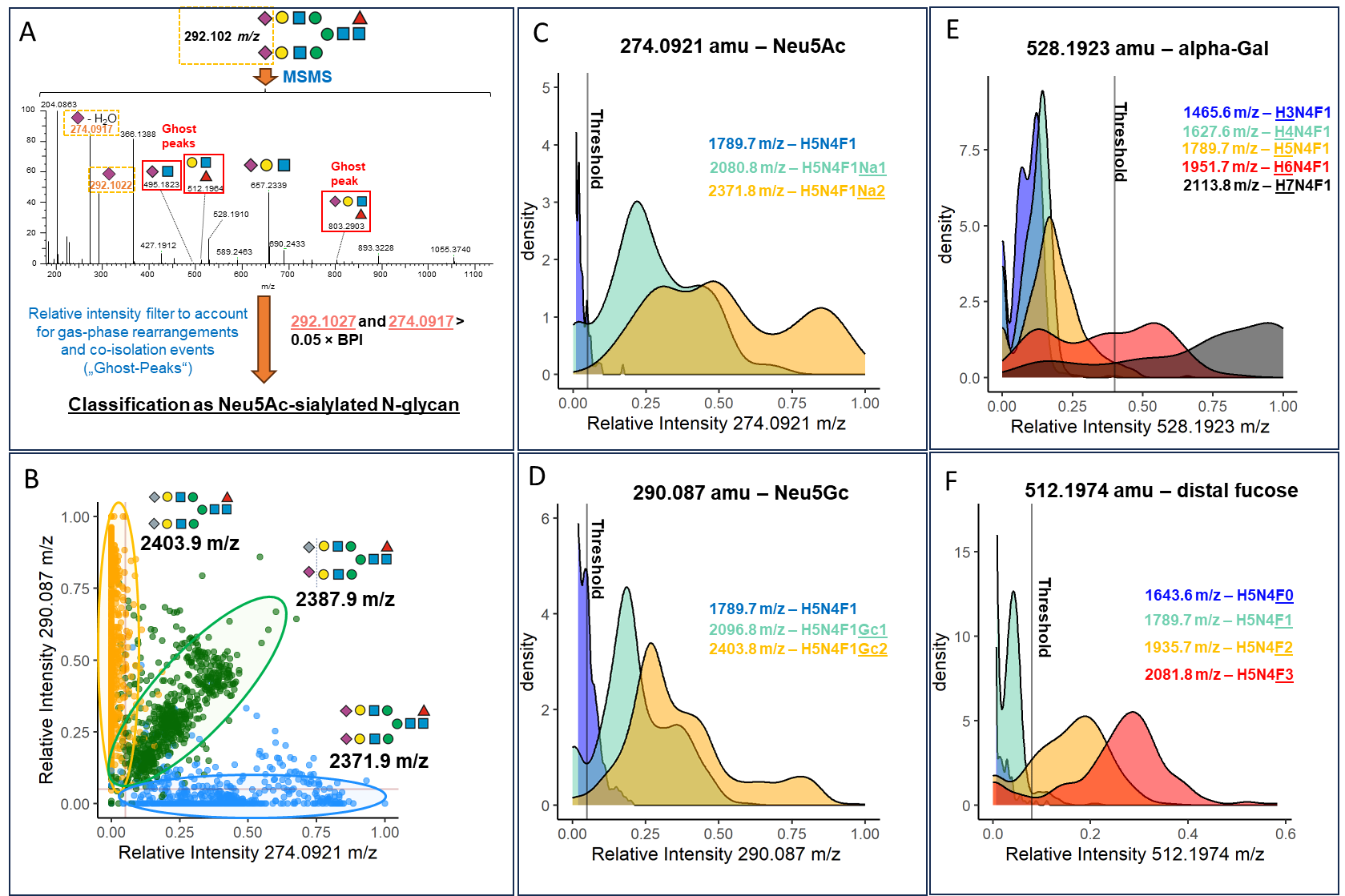

**SI Figure 6. Empiric determination of the cut off values for the extended SNOG-scores (eSNOG). (A)** Representative MS/MS spectrum of a Neu5Ac-sialylated N-glycan with the composition Hex_5_HexNAc_4_Fuc_1_Neu5Ac_2_. Red boxed fragment ions are the result of gas-phase rearrangements of Neu5Ac or fucose residues. The simultaneous presence of both the 292.1 amu and 274.1 amu fragment ions are used to classify precursors as “Neu5Ac-sialylated”. **(B)** Relative Intensities of the Neu5Gc-specific fragment ion (290.1 amu), and the Neu5Ac-specific fragment ion (274.1 amu) derived from three different N-glycan precursors, i.e., Hex_5_HexNAc_4_Fuc_1_Neu5Gc_2_ (orange dots), Hex_5_HexNAc_4_Fuc_1_Neu5Ac_1_Neu5Gc_1_ (green dots), and Hex_5_HexNAc_4_Fuc_1_Neu5Ac_2_ (blue dots) across all samples. **(C)** Density plot of the relative intensity of the 274 .1 amu fragment ion diagnostic for Neu5Ac-sialylated N-glycans for three different N-glycan compositions, i.e., Hex_5_HexNAc_4_Fuc_1_ (blue), Hex_5_HexNAc_4_Fuc_1_Neu5Ac_1_ (turquoise), and Hex_5_HexNAc_4_Fuc_1_Neu5Ac_2_ (orange) across all tissues. The plot is cut-off at a density value of 5. An eSNOG_274_-score of 0.05 efficiently discriminated Neu5Ac-sialylated N-glycans. **(D**) Density plot of the relative intensity of the 290.1 amu fragment ion, diagnostic for Neu5Gc-sialylated N-glycans, for three different N-glycan compositions, i.e., Hex_5_HexNAc_4_Fuc_1_ (blue), Hex_5_HexNAc_4_Fuc_1_Neu5Gc_1_ (turquoise), and Hex_5_HexNAc_4_Fuc_1_Neu5Gc_2_ (orange) across all tissues. The plot is cut-off at a density value of 6. An eSNOG_290_-score of 0.05 efficiently discriminated Neu5Gc-sialylated N-glycans. **(E)** Density plot of the relative intensity of the 528.2 amu fragment ion for five different compositions (i.e., Hex_3_HexNAc_4_Fuc_1_ (blue), Hex_4_HexNAc_4_Fuc_1_ (turquoise), Hex_5_HexNAc_4_Fuc_1_ (orange), Hex_6_HexNAc_4_Fuc_1_ (red), Hex_7_NAc_4_Fuc_1_ (black), and Hex_7_HexNAc_4_Fuc_1_ across all tissues. With some exceptions, Hex_6_HexNAc_4_Fuc_1_ (red) and Hex_7_NAc_4_Fuc_1_ (black) are considered to contain alpha galactose. An eSNOG_528_-score of 0.35 efficiently discriminated alpha-galactosylated N-glycans. **(F)** Density plot of the relative intensity of the 512.2 amu fragment ion, diagnostic for antenna fucosylation, for four different compositions (i.e., Hex_5_HexNAc_4_ (blue), Hex_5_HexNAc_4_Fuc_1_ (turquoise), Hex_5_HexNAc_4_Fuc_2_ (orange), and Hex_5_HexNAc_4_Fuc_3_ (red) across all tissues. With some exceptions, the composition Hex_5_HexNAc_4_Fuc_1_ (turquoise) is considered to be core fucosylated. An eSNOG_512_-score of 0.075 efficiently discriminated antenna-fucosylated N-glycans and accounts for the potential gas-phase rearrangement of core fucose residues.

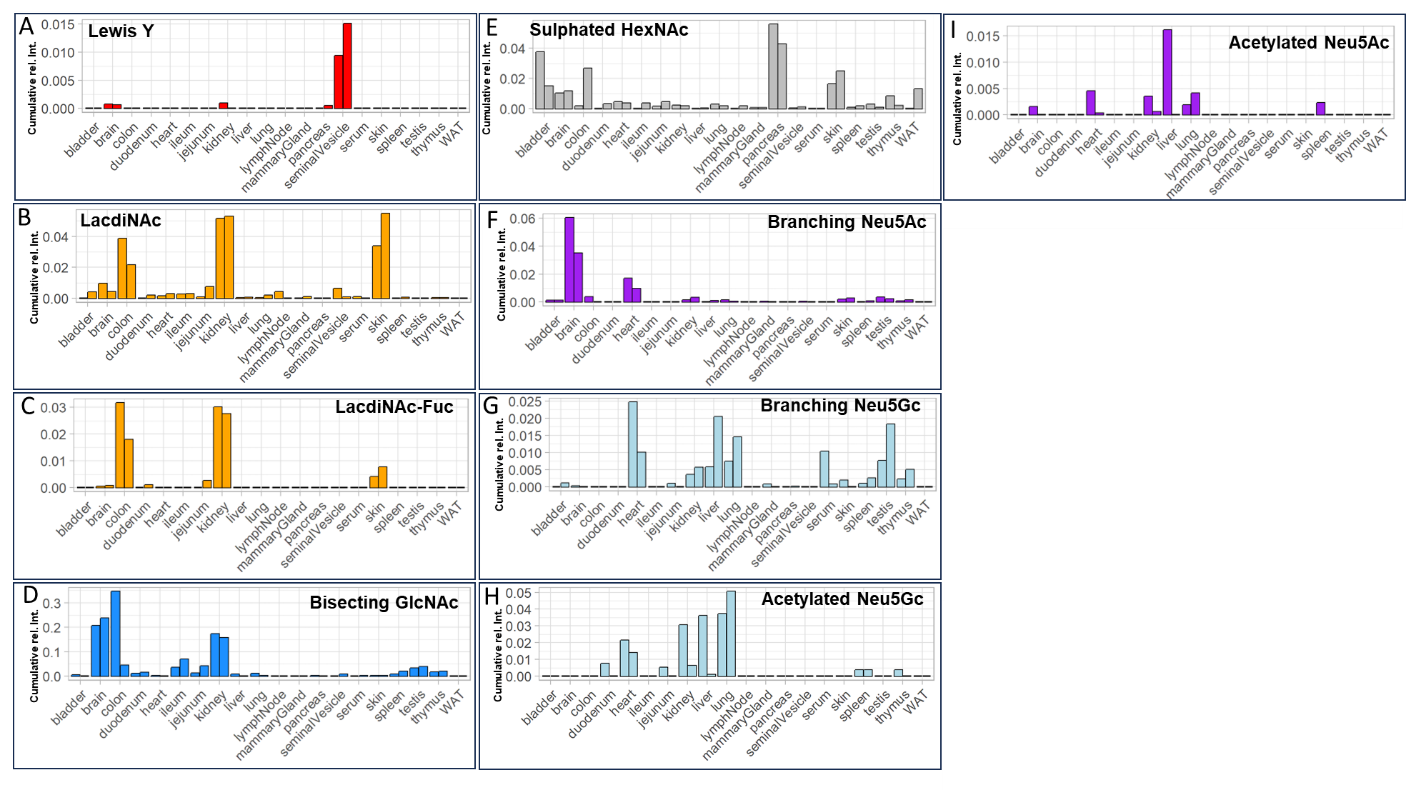

**SI Figure 7.** **Tissue specific expression of specific N-glycan modifications. (A)** Lewis Y epitope. Comparative analysis of N-glycans carrying the Lewis Y epitope across tissues. **(B)** LacdiNAc modification. Comparative analysis of N-glycans carrying the LacdiNAc modification across tissues. **(C**) Fucosylated LacdiNAc modification. Comparative analysis of N-glycans carrying the fucosylated LacdiNAc modification across tissues. **(D**) Bisecting GlcNAc. Comparative analysis of N-glycans carrying a bisecting GlcNAc across tissues. **(E)** Sulfated HexNAc. Comparative analysis of N-glycans carrying sulfated GlcNAc across tissues. **(F**) Branching Neu5Ac. Comparative analysis of N-glycans carrying branching Neu5Ac across tissues. (G) Branching Neu5Gc. Comparative analysis of N-glycans carrying branching Neu5Gc across tissues. **(H)** Acetylated Neu5Gc. Comparative analysis of N-glycans carrying acetylated Neu5Gc across tissues. **(I)** Acetylated Neu5Ac. Comparative analysis of N-glycans carrying acetylated Neu5Ac across tissues. All values are normalized to the total glyco-TIC of the respective tissue. “WAT” - white adipose tissue.

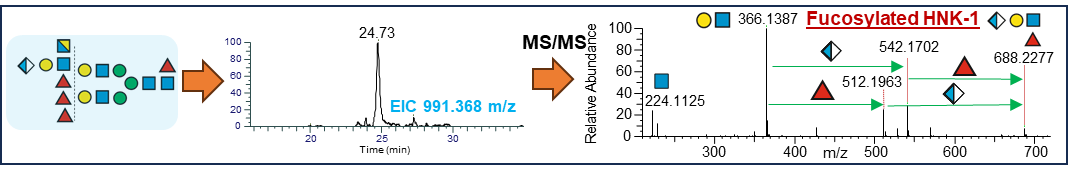

**SI Figure 8. MS/MS supported identification of a N-glycan with the composition Hex_6_HexNAc_6_Fuc_4_GlcA_1_ carrying a fucosylated HNK-1 epitope in kidney.**

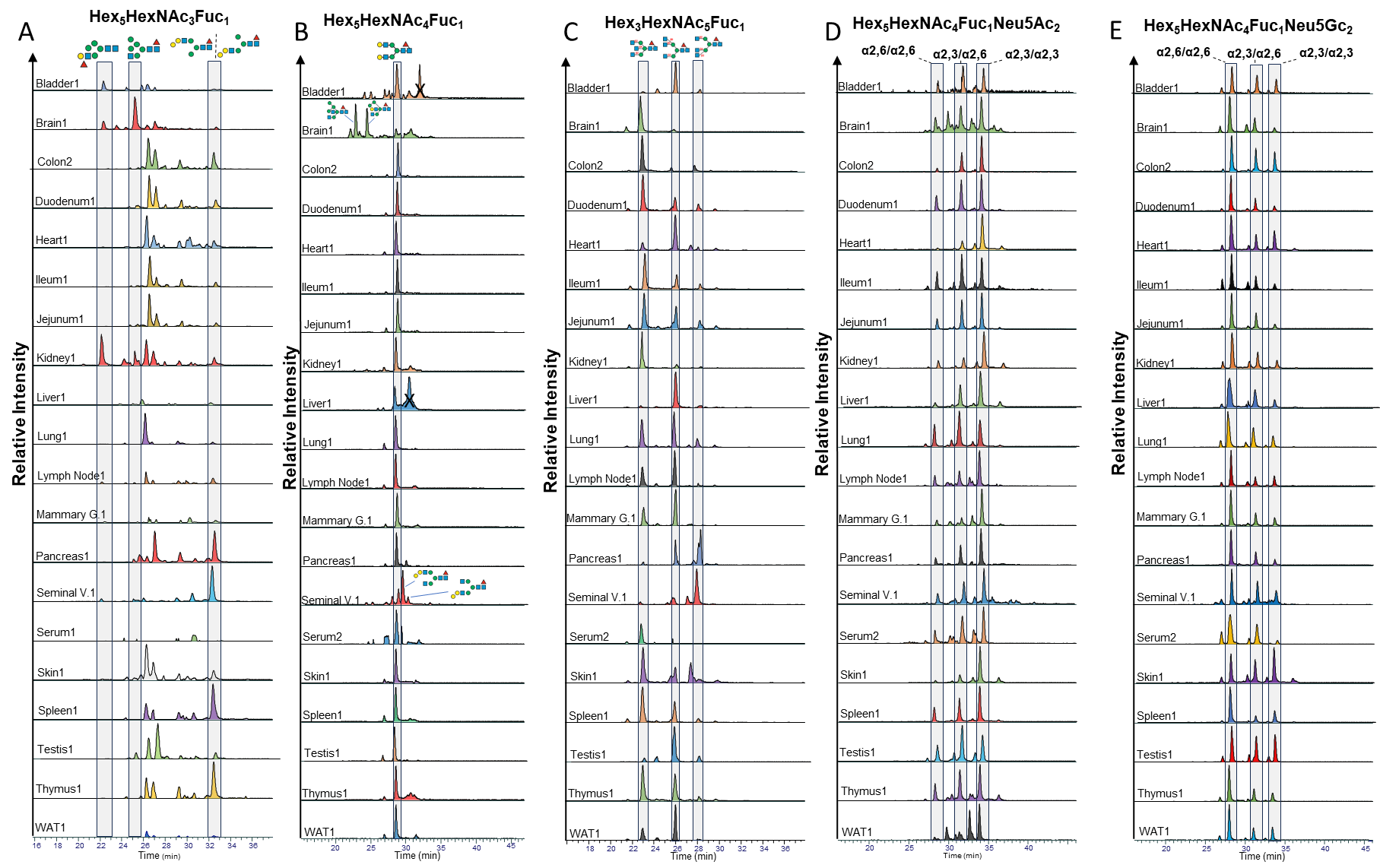

**SI Figure 9.** **Profiling the isomeric complexity of the murine N-glycome**. The highly isomer-selective stationary phase porous graphitic carbon (PGC) was used to separate even closely related isomers. Already existing N-glycan retention libraries combined with MS/MS data were used to identify the exact structures of the respective isomers. All retention times were normalized to the retention time of the ubiquitous Man5 N-glycan. **(A)** shows the elution profile for the composition Hex_5_HexNAc_3_Fuc_1_ across all analyzed tissues. **(B)** shows the elution profile for the composition Hex_5_HexNAc_4_Fuc_1_ across all analyzed tissues. **(C**) shows the elution profile for the composition Hex_3_HexNAc_5_Fuc_1_ across all analyzed tissues. (D) shows the elution profile for the composition Hex_5_HexNAc_4_Fuc_1_Neu5Ac_2_ across all analyzed tissues. **(E)** shows the elution profile for the composition Hex_5_HexNAc_4_Fuc_1_Neu5Gc_2_ across all analyzed tissues.

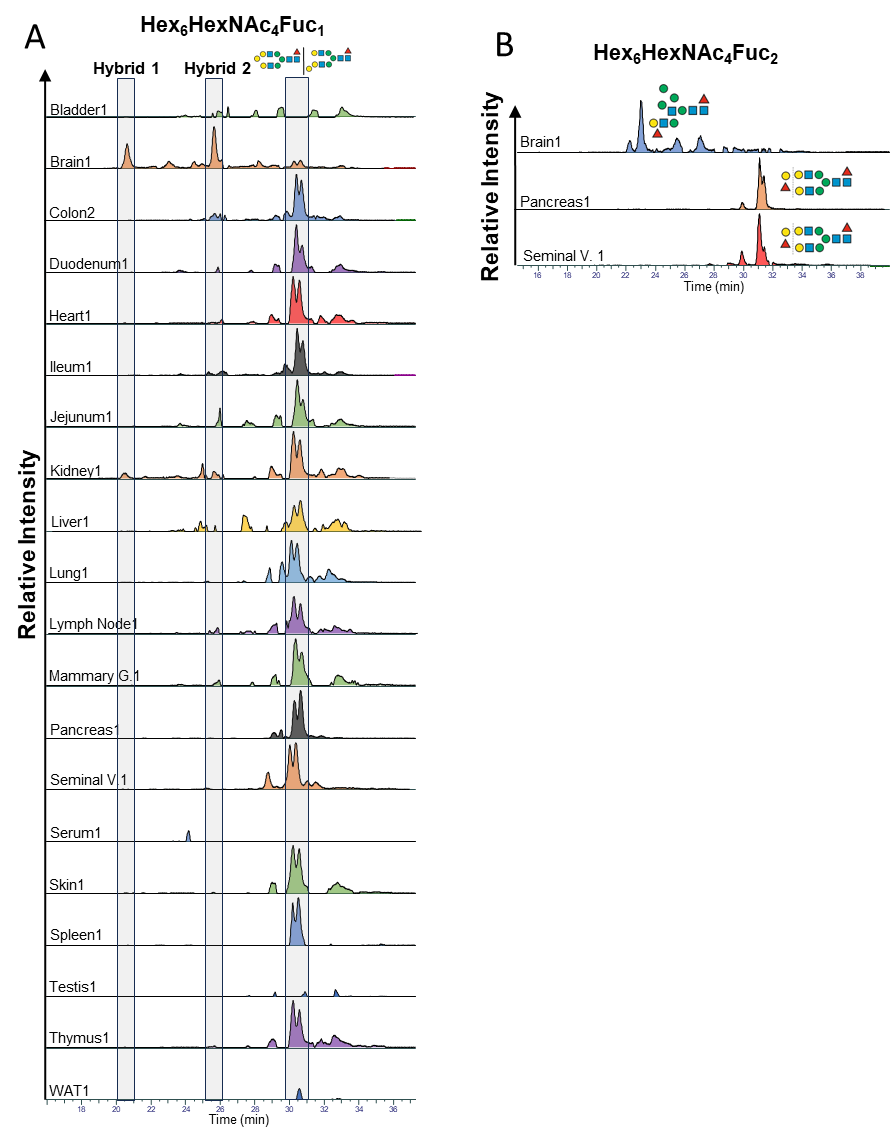

**SI Figure 109.** **Profiling the isomeric complexity of the murine N-glycome**. **The highly isomer-selective stationary phase porous graphitic carbon (PGC) was used to separate even closely related isomers. Already existing N-glycan retention libraries combined with MS/MS data were used to identify the exact structures of the respective isomers. All retention times were normalized to the retention time of the ubiquitous Man5 N-glycan. (A) shows the elution profile for the composition Hex_6_HexNAc_3_Fuc_1_ across all analyzed tissues. (B) shows the elution profile for the composition Hex_6_HexNAc_4_Fuc_2_ across brain, pancreas, and seminal vesicle.** “WAT” - white adipose tissue.

**Supplementary Tables**

**SI Table 1. Diagnostic fragment ions and associated eSNOG-score cutoffs used for sub-structural profiling of N-glycans.**

| **Modification** | **Diagnostic fragment ions [H^+^] [amu]** | **eSNOG-score cutoff** |
| --- | --- | --- |
| Neu5Ac | 292.1072 & 274.0921 | 0.025 & 0.05 |
| Neu5Gc | 308.0976 & 290.087 | 0.025 & 0.05 |
| branching Neu5Ac | 495.1821 | 0.025 |
| branching Neu5Gc | 511.177 | 0.025 |
| Neu5Ac-associated di-sialyl Lewis C | 495.1821 & 948.3303 | 0.025 & 0.005 |
| Neu5Gc-associated di-sialyl Lewis C | 511.177 & 980.3201 | 0.025 & 0.005 |
| Acetylated Neu5Ac | 334.1133 & 316.1027 | 0.005 & 0.005 |
| Acetylated Neu5Gc | 350.1082 & 332.0976 | 0.005 & 0.005 |
| Antennary fucose | 512.1974 | 0.075 |
| Alpha-galactose | 528.1923 | 0.35 |
| HNK-1 epitope | 542.1716 | 0.05 |
| Bisecting GlcNAc | 792.3234 | 0.025 |
| LacdiNAc | 407.1661 | 0.04 |
| Fucosylated LacdiNAc | 407.1661 & 553.224 | 0.04 & 0.01 |
| Lewis Y | 512.1974 & 658.2553 | 0.2 & 0.07 |
| Sulphated HexNAc | 284.0435 | 0.01 |
| Reduced N-glycan | 224.1118 | 0.03 |
