## Supplementary figures and images for "Non-targeted isomer-sensitive N-glycome analysis reveals new layers of organ-specific diversity in mice"

### Precursor-independent MS/MS-based profiling for fucose across all tissues

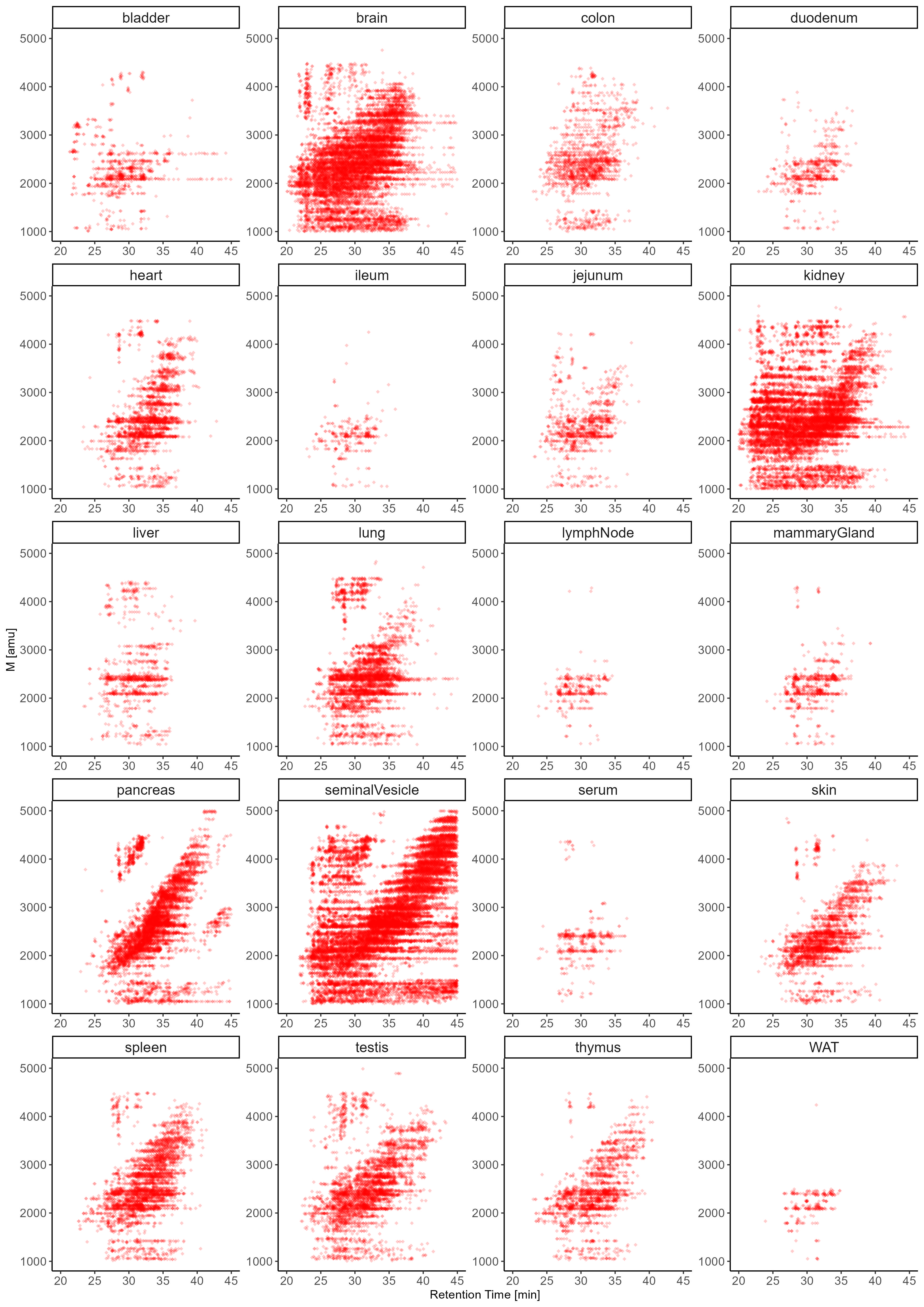

### Precursor-independent MS/MS-based profiling for Neu5Ac across all tissues

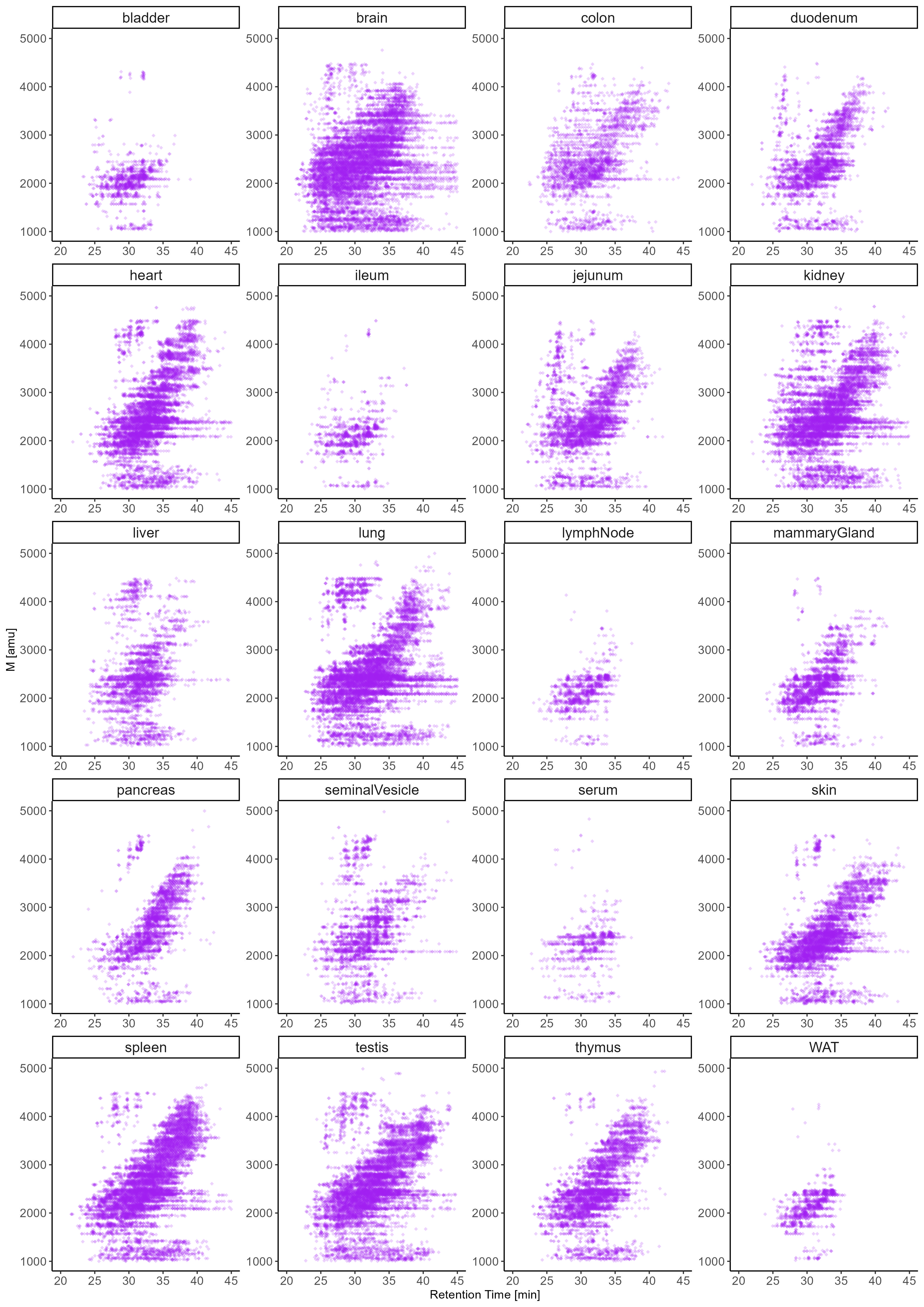

### Precursor-independent MS/MS-based profiling for Neu5Gc across all tissues

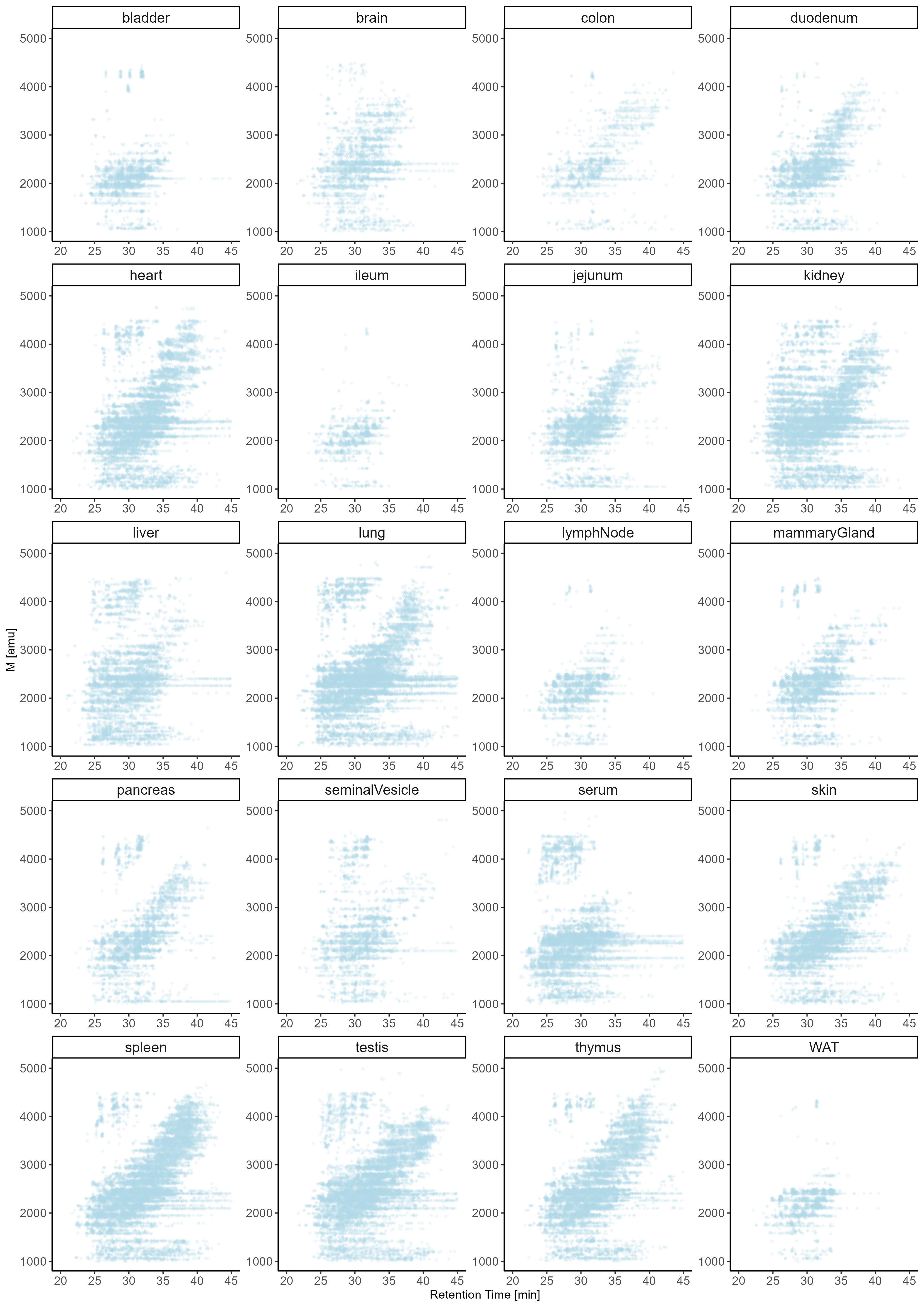

### Semi-quantitative histograms of Antennary-fucose-associated eSNOG-filtered LC-MS data of all samples

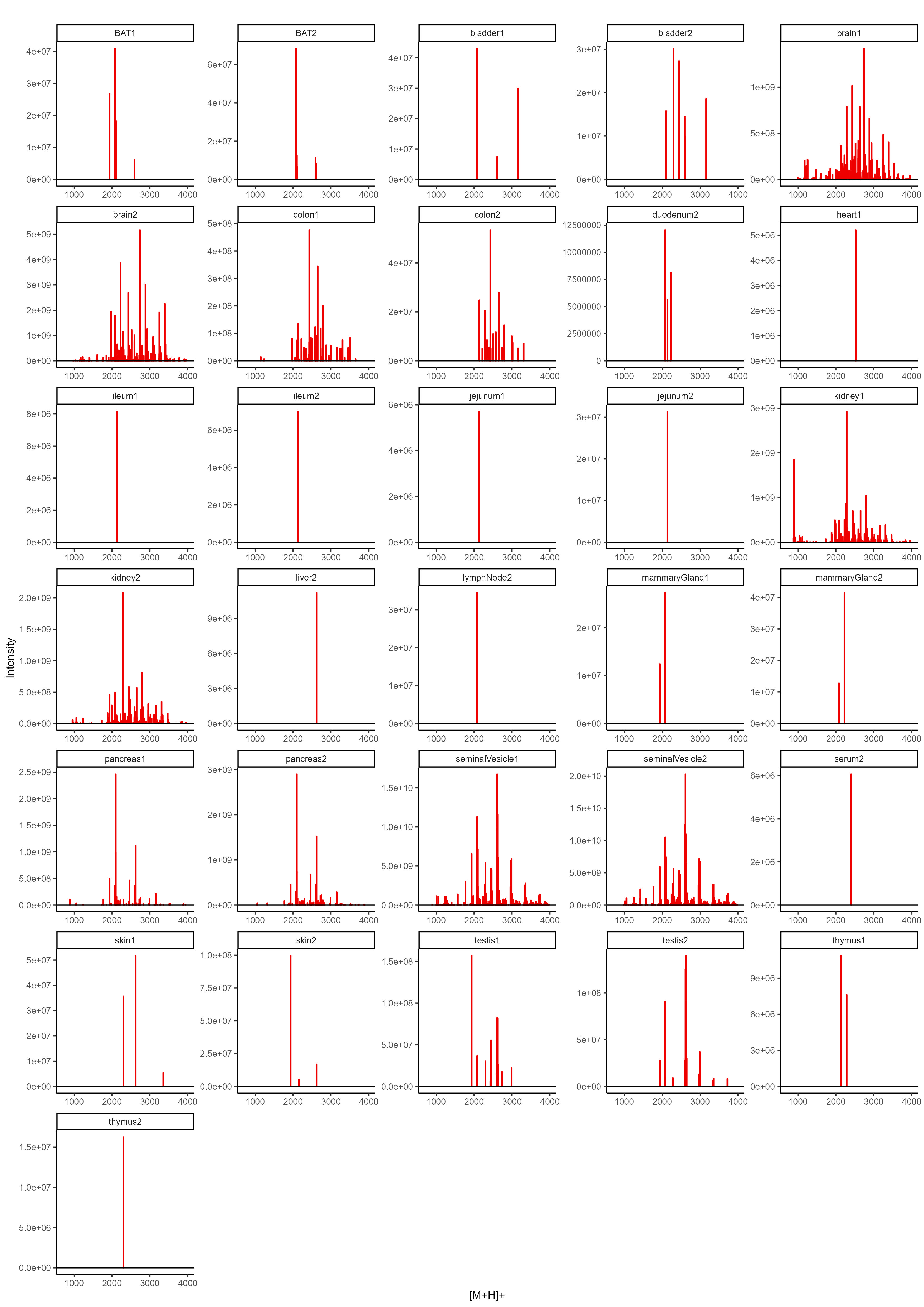

### Semi-quantitative histograms of Neu5Ac-associated eSNOG-filtered LC-MS data of all samples

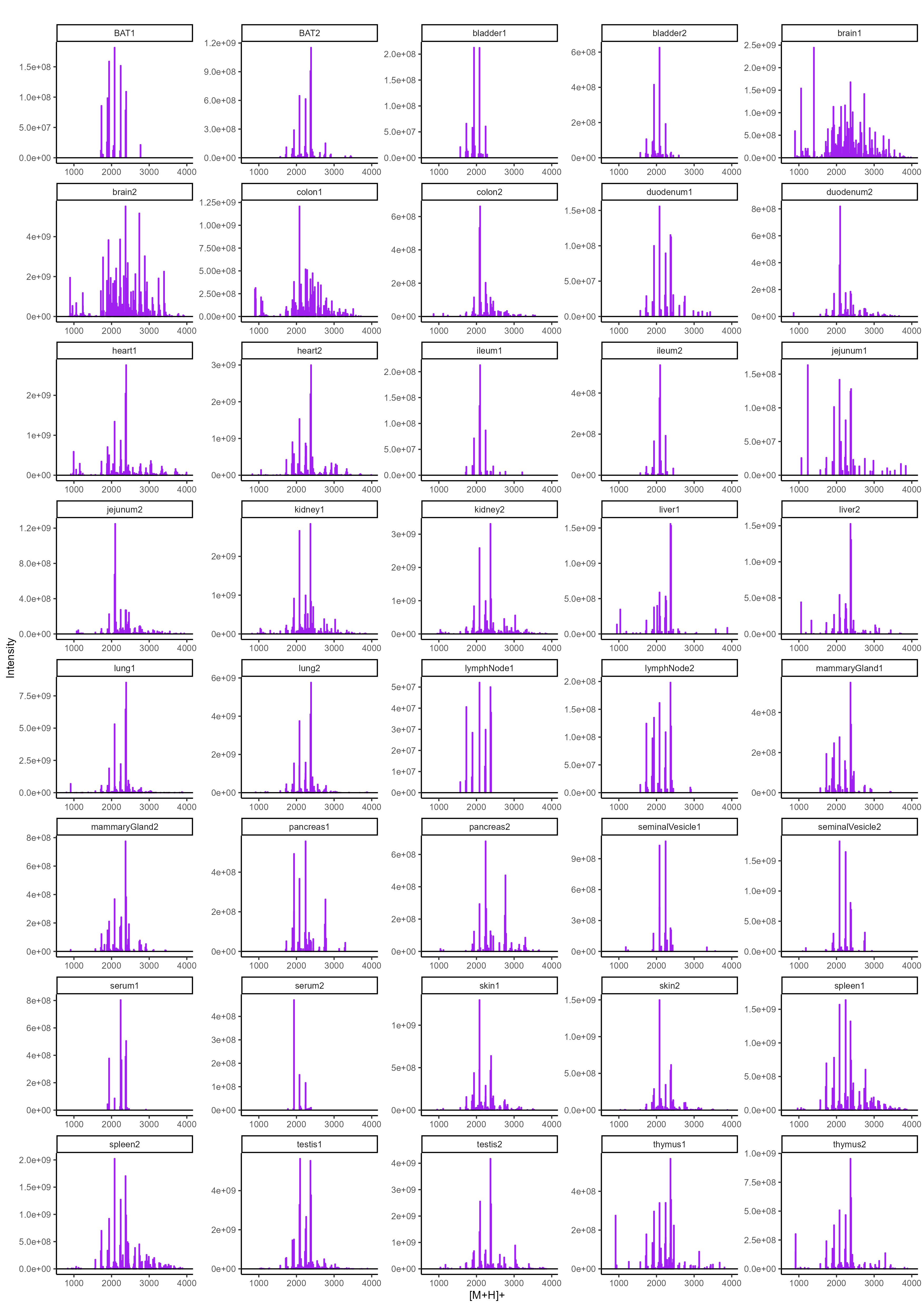

### Semi-quantitative histograms of Neu5Gc-associated eSNOG-filtered LC-MS data of all samples

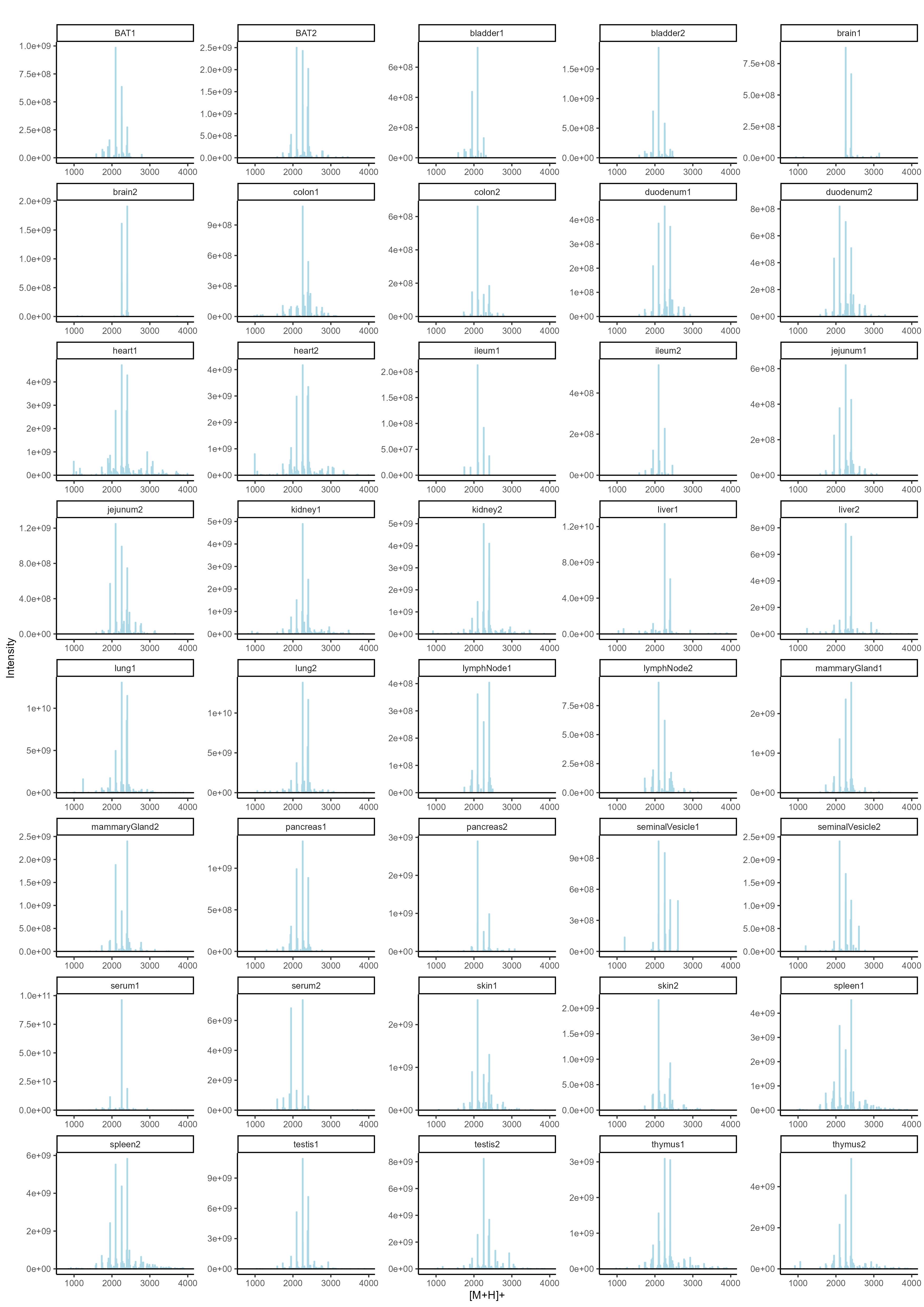

### Semi-quantitative histograms of SNOG-filtered LC-MS data of all samples

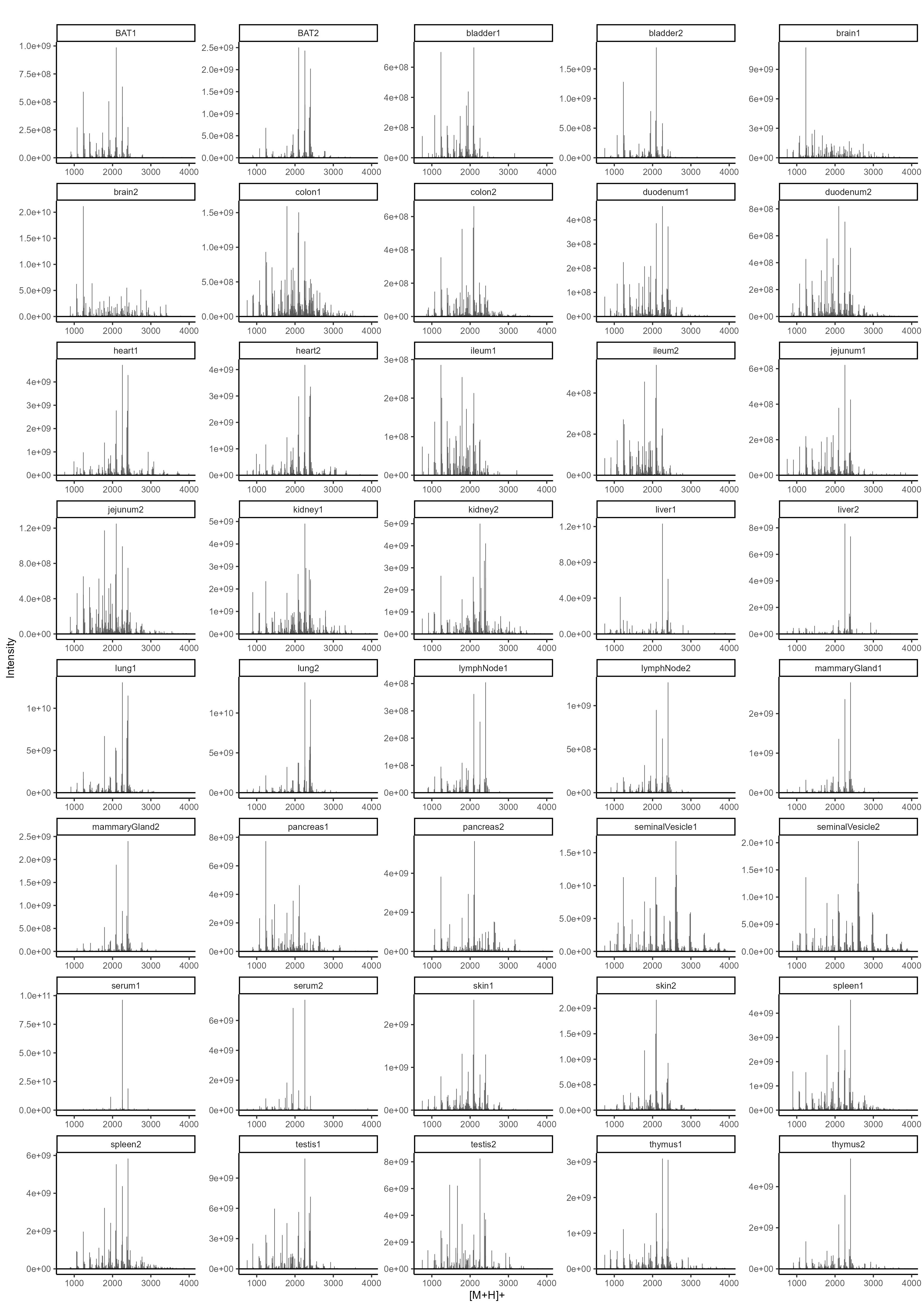
